# Torso-like is a component of the hemolymph and regulates the insulin signalling pathway in *Drosophila*

**DOI:** 10.1101/264325

**Authors:** Michelle A. Henstridge, Lucinda Aulsebrook, Takashi Koyama, Travis K. Johnson, James C. Whisstock, Tony Tiganis, Christen K. Mirth, Coral G. Warr

**Affiliations:** School of Biological Sciences, Monash University, Clayton, Victoria 3800, Australia.; Instituto Gulbenkian de Ciência, Oeiras, Portugal; Department of Biochemistry and Molecular Biology, Monash University, Clayton, Victoria 3800, Australia.; Monash Biomedicine Discovery Institute, Monash University, Clayton, Victoria 3800, Australia.

**Keywords:** Torso-like, Insulin Signalling, Body size, Developmental timing, *Drosophila melanogaster*

## Abstract

In *Drosophila* key developmental transitions are governed by the steroid hormone ecdysone. A number of neuropeptide-activated signalling pathways control ecdysone production in response to environmental signals, including the insulin signalling pathway, which regulates ecdysone production in response to nutrition. Here, we find that the Membrane Attack Complex/Perforin-like protein Torso-like, best characterised for its role in activating the Torso receptor tyrosine kinase in early embryo patterning, also regulates the insulin signalling pathway in *Drosophila*. We previously reported that the small body size and developmental delay phenotypes of *torso-like* null mutants resemble those observed when insulin signalling is reduced. Here we report that, in addition to growth defects, *torso-like* mutants also display metabolic and nutritional plasticity phenotypes characteristic of mutants with impaired insulin signalling. We further find that in the absence of *torso-like* the expression of insulin-like peptides is increased, as is their accumulation in the insulin-producing cells. Finally, we show that Torso-like is a component of the hemolymph and that it is required in the prothoracic gland to control developmental timing and body size. Taken together, our data suggest that the secretion of Torso-like from the prothoracic gland influences the activity of insulin signalling throughout the body in *Drosophila*.

**ARTICLE SUMMARY:** In many animals distinct developmental transitions are crucial for the coordinated progression from the juvenile stage to adulthood. In *Drosophila*, the transition from an immature larva into a reproductively mature adult is controlled by the steroid hormone ecdysone. Several neuropeptide-activated signalling pathways, including the insulin signalling pathway, regulate ecdysone production in response to environmental cues. Here we find that the perforin-like protein Torso-like regulates the insulin signalling pathway. We show that Torso-like is secreted into circulation where it acts to influence insulin-like peptide activity, revealing a novel mechanism for the regulation of insulin signalling in *Drosophila.*

## INTRODUCTION

In holometabolous insects such as *Drosophila*, pulses of the steroid hormone ecdysone regulate the timing of developmental transitions, including the metamorphic transition, and hence growth duration (Henrich *et al.* 1999; Mirth & Riddiford 2007). During the larval and pupal stages of development ecdysone is produced and released by the major endocrine organ, the prothoracic gland (PG), in response to multiple environmental and developmental stimuli (for review see Danielsen *et al.* 2013). Accordingly, there are many complex cellular signalling pathways involved in coordinating the responses to these signals. A well-studied example is prothoracicotropic hormone (PTTH), a brain-derived neuropeptide that regulates the production and release of ecdysone in response to developmental cues (McBrayer *et al.* 2007). Prior to each larval moult, PTTH is secreted and activates the Torso (Tor) receptor tyrosine kinase (RTK) in the PG, which signals via the Ras/mitogen-activated protein kinase pathway to upregulate a set of ecdysone biosynthesis genes (Rewitz *et al.* 2009). Ablation of the PTTH-neurons, or loss of function mutations in *tor*, prolongs the growth period between each developmental transition and results in an overall increase in body size (McBrayer *et al.* 2007; Rewitz *et al.* 2009).

Another critical signalling pathway that regulates ecdysone production is the evolutionarily conserved insulin signalling pathway, which acts in the PG to regulate ecdysone biosynthesis in response to nutrition (Caldwell *et al.* 2005; Colombani *et al.* 2005; Mirth *et al.* 2005). In particular, this pathway regulates larval growth rate and the timing of a developmental checkpoint known as critical weight, thereby controlling the timing of the onset of metamorphosis (Mirth *et al.* 2005; Koyama *et al.* 2014). In *Drosophila,* the insulin-like receptor (InR) is activated by a family of insulin-like peptides (dILPs) (Brogiolo *et al.* 2001). A subset of these (dILPs 2, 3 and 5) are expressed in neurons that innervate the corpora cardiaca, a group of cells neighbouring the PG, and ablation of these neurons causes developmental delays and decreased body size (Rulifson *et al.* 2002).

We and others recently reported that mutant alleles of the Membrane Attack Complex/ Perforin-like (MACPF) protein Torso-like (Tsl) exhibit a developmental delay phenotype indicative of defects in ecdysone production (Grillo *et al.* 2012; Johnson *et al.* 2013). Tsl is best known for its role in embryonic patterning, where it functions upstream of the Tor receptor to control its activity for patterning the embryo termini (Stevens *et al.* 1990; Savant-Bhonsale & Montell 1993; Martin *et al.* 1994). Unexpectedly, however, we provided evidence that Tsl does not act similarly with Tor in the PG to control developmental transitions and body size. Specifically, *tsl* and *tor* have opposing effects on body size, and the developmental delay phenotype observed in *tsl;tor* double mutants is strikingly enhanced compared to either mutation alone, suggesting an additive rather than epistatic interaction (Johnson *et al.* 2013).

The growth defects of *tsl* mutants more closely resemble those observed when insulin signalling is reduced, however, Tsl has not previously been implicated in the insulin signalling pathway. Furthermore, it is not clear whether the growth and developmental timing phenotypes of *tsl* null mutants are due to a role for Tsl in the PG. Here, we report that in addition to growth defects, *tsl* mutants also display several other physiological and biochemical characteristics of impaired insulin signalling. We further show that Tsl is required in the PG to control developmental timing and body size, and that it influences the expression of dILPs and their accumulation in the insulin-producing cells (IPCs). Finally, we show that Tsl is present in the larval hemolymph, strongly supporting the idea that Tsl is secreted from the PG into circulation where it acts to regulate systemic insulin signalling.

## MATERIALS AND METHODS

### *Drosophila* stocks

The following stocks were used: *w^1118^* (BL5905), *chico^1^* (BL10738), Df(2L)BSC143 (*chico^df^*; (BL9503) a chromosomal deficiency that deletes the *chico* coding region), *InR^A1325D^* (BL8263; a constitutively active form of InR), and *dilp2-3^∆^,dilp5^3^* (BL30889) from the Bloomington *Drosophila* stock centre; *c7-Gal4*, obtained from FlyView (Janning, 1997); *tsl*^∆^, a null mutant of *tsl* (Johnson *et al.* 2013); *phm-*Gal4 (ch2) and UAS-*dicerII; phm-*Gal4-22, gifts from Michael O’Connor, University of Minnesota, Minneapolis (Ono *et al.* 2006); and UAS-*tsl^RNAi^* and *gHA:tsl*, gifts from Jordi Casanova, IRB Barcelona (Jimenez *et al.* 2002; Furriols *et al.* 2007). All flies were maintained at 25°C on fly media containing, per litre: 7.14 g potassium tartrate, 0.45 g calcium chloride, 4.76 g agar, 10.71g yeast, 47.62g dextrose, 23.81g raw sugar, 59.52g semolina, 7.14mL Nipagen (10% in ethanol) and 3.57mL propionic acid.

### Torso-like constructs and generation of transgenic lines

To generate the UAS-*Tsl:HA* and UAS-*Tsl:RFP* constructs, the open reading frame of *tsl* followed by a short linker encoding the peptide SAGSAS and either three tandem HA epitopes (for UAS-*Tsl:HA*) or the open reading frame for RFP (for UAS-*Tsl:RFP*) was synthesised and subcloned (Genscript) into pUASTattB via BglII and XhoI sites. To generate the *phm:Tsl* construct, a 1.1 kb fragment of the *phm* promoter region (from Ono *et al.* (2006)) was first cloned from gDNA (F – 5’-CTG CAG TGA TGC GCT GCT CCT TTG T-3’, R – 5’-AGA TCT CAC TTT CGA TTT CCT CCT GC-3’) into the pGEM-T Easy vector (Promega), before being sequenced and subcloned into pUASTattB-*Tsl:eGFP* (Johnson *et al.* 2017) via PstI and BglII sites to delete the UAS sequence. Transgenic lines were made (BestGene) via ΦC31 integrase mediated transformation (Bischof *et al.* 2007) using the ZH-51CE attP-landing site.

### Developmental timing and body size analysis

Twenty-four hours after a 4 h lay on apple juice agar supplemented with yeast paste, first-instar larvae were sorted by GFP (on a balancer chromosome) into 8–10 groups of 15 or 20 individuals (depending upon experiment) per genotype. Larvae were placed into vials containing fly media (see recipe above) and scored every 8 h for the time taken to reach pupariation. Following their eclosion, adult flies were sorted by sex and weighed in groups on a microbalance (Mettler Toledo).

### Nutritional plasticity

Twenty-four hours after a 4 h lay on apple juice agar supplemented with yeast paste, first-instar larvae were sorted by GFP (on a balancer chromosome) and placed into vials containing either standard fly media or one of three low nutrient diets (either 50, 25 or 10% nutrients of standard media). These diets were made by diluting standard fly media with 0.5% non-nutritional agar to the appropriate concentration. Adult flies were collected within 24 h of eclosion, sorted and weighed in groups on a microbalance (Mettler Toledo). For each genotype, 10 replicates of 15 larvae were raised on each food type. Because size increases exponentially with increasing nutritional quality, male and female weight data was log10 transformed and analysed by fitting the log10 transformed weights with linear models, using food concentration and genotype as fixed effects, in R-studio. Significant interactions between food concentration and genotype on body weight indicates that the two genotypes show significant differences in nutritional plasticity for body weight.

### Quantification of food intake

Early feeding third-instar larvae were transferred to fresh dyed food (4.5% blue food dye) and allowed to feed for 1 h. After feeding, larvae were removed from food using 20% sucrose solution, washed in distilled water and dried. Replicates of 10 larvae were homogenised in 80µl of cold methanol and centrifuged for 10 min at 4°C. 60µl of supernatant from each sample was analysed in a spectrophotometer at 600nm. As standards, a two-fold dilution series of food dye, using a starting concentration of 4µl dye/ml methanol was used. Five biological replicates were analysed per genotype.

### Hemolymph glucose and trehalose measurements

Hemolymph was pooled from 15 wandering third-instar larvae to obtain duplicate samples of 1µl for assay. Five biological replicates were performed per genotype. Glucose was measured by adding 99µl of Thermo Infinity Glucose Reagent (Thermo Scientific) to each sample and processing as per the manufacturer’s instructions. Trehalose was measured using the same reagent after digestion to glucose using trehalase, with a ten-fold dilution due to higher levels of trehalose compared to glucose. For trehalose digestion 1µl of hemolymph was incubated in 25µl of 0.25M sodium carbonate at 95°C for 2 h, cooled to room temperature, and 8µl of 1M acetic acid and 66µl of 0.25M sodium acetate (pH 5.2) were added to make digestion buffer. 1µl of porcine trehalase (Sigma, T8778) was added to 40µl of this mixture and incubated at 37°C overnight. The resulting glucose was analysed using 10µl of reaction and 90µl Thermo Infinity Glucose Reagent as above. Glucose and trehalose standards were treated together with samples to quantify sugar levels in hemolymph.

### Whole body triglyceride measurements

Triglycerides were quantified in whole wandering third-instar larvae as per Musselman *et al.* (2011). Ten larvae were homogenised in phosphate buffered saline (PBS) + 0.1% Tween and then diluted 1:100 with PBS. Samples were heated for 5 min at 65°C to inactivate lipases and 2µl of each sample was mixed with 198µl Thermo Infinity Triglyceride Reagent (Thermo Scientific) and processed as per the manufacturer’s instructions. The absorbance of samples at 500nm was used as a relative measure of triglyceride content and was normalised to larval weight. Five biological replicates of ten larvae were analysed in duplicate for each genotype.

### Immunoblotting

Hemolymph was extracted from approximately 80 wandering third instar larvae on ice. Following centrifugation at 16,000g for 5 min at 4°C, supernatant was heat-inactivated at 60°C for 10 min, re-spun and the remaining supernatant was combined with 1 mM DTT, 10 mM NaF and complete EDTA-free protease inhibitor cocktail (Roche). For pAkt blots, five third instar larvae were homogenised in 80µl of lysis buffer (50mM Tris-HCl (pH 7.5), 150mM NaCl, 2.5mM EDTA, 0.2% Triton X, 5% glycerol, complete EDTA-free protease inhibitor cocktail (Roche)) and spun at 500g for 5 min at 4°C. Reducing buffer (containing 6M urea for Tsl immunoblots) was added to all samples before boiling and separation by SDS-PAGE (any kDa TGX, Biorad) followed by transfer onto an Immobilon-P membrane (Millipore). Membranes were probed with either 1:1,000 anti-HA (Roche, 12CA5), 1:1,000 anti-phosphorylated *Drosophila* Akt (Cell Signalling, 4054S), or 1:1,000,000 anti-α-tubulin (Sigma, B-5-1-2), washed and incubated with HRP-conjugated secondary antibody (1:10,000, Southern Biotech). Immunoblots were developed using ECL prime (GE healthcare) and imaged using a chemiluminescence detector (Vilber Lourmat). pAkt blot images were quantified using ImageJ and differences between genotypes determined by unpaired *t*-tests from five biological replicates.

### Immunostaining and fluorescence quantification

Newly molted L3 larvae were collected and allowed to age on standard media for 24 h before brains were dissected and fixed in phosphate buffered saline (PBS) containing 4% paraformaldehyde for 40 min. Tissues were extensively washed in PBS containing 0.3% Triton X-100 (PTx) and then blocked for 1 h in PBT containing 2% normal goat serum (Sigma-Aldrich). Primary antibodies (rat anti-dILP2 and rabbit anti-dILP5; gifts from Dr. Pierre Léopold (Geminard *et al.* 2009)) were diluted to 1:800 in fresh block solution and incubated overnight at 4°C. After extensive PTx washes, secondary antibodies (anti-rat Alexa 488, and anti-rabbit Alexa 568 conjugated; Molecular Probes) were diluted to 1:500 and incubated over night at 4ºC. Brains were mounted in Fluoromount-G mounting medium (Southern Biotech) and Z series of the insulin-producing cells (IPCs) were obtained using a spinning disk confocal microscope (Olympus CV1000), maintaining a 1µm step size and identical imaging settings across all genotypes. ImageJ software was used to generate maximum projection images of the Z stacks and to quantify total fluorescent intensity across the IPCs. This was achieved by drawing an area of interest around each group of IPCs and calculating the raw grey scale values in this region of interest. The total fluorescence was normalised to IPC area to account for the size discrepancy between genotypes.

### Insulin-like peptide gene transcript quantification

For each biological replicate, 10–15 third instar larvae (anterior end only) or 20–25 dissected third instar larval brains were snap frozen before RNA was extracted using TRIsure reagent (Bioline) and treated with DNAse (Promega). Complementary DNA was synthesised using Tetro reverse transcriptase (Bioline) by priming either 5µg (for anterior ends) or 1 µg (for dissected brains) of RNA with oligo-dT and random hexamers. Quantitative PCRs were performed in triplicate on a Light Cycler 480 (Roche) using SensiMix Sybr (Bioline) and primers specific for *dilp2* (F – 5’-ACG AGG TGC TGA GTA TGG TGT GCG-3’, R – 5’-CAC TTC GCA GCG GTT CCG ATA TCG-3’)*, dilp5* (F – 5’-TGT TCG CCA AAC GAG GCA CCT TGG-3’, R – 5’-CAC GAT TTG CGG CAA CAG GAG TCG-3’) and *Rp49* (F – 5’-GCC GCT TCA AGG GAC AGT ATC T-3’, R – 5’-AAA CGC GGT TCT GCA TGA G-3’). Fold changes relative to *Rp49* were determined using the delta CT method and means and standard errors calculated from 3–5 biological replicates per genotype.

### Data and reagent availability statement

Data and reagents are available upon request.

## RESULTS

In addition to its role in regulating growth and developmental timing, the insulin signalling pathway is also critical for regulating glucose and lipid metabolism in *Drosophila* (for review see Garofalo, 2002). Thus, in addition to growth defects, mutants with reduced insulin signalling are unable to regulate their blood-sugar levels (Rulifson *et al.* 2002). This results in increased levels of glucose in the hemolymph (but not of trehalose - a glucose disaccharide that is synthesised from intracellular glucose in the fat body and secreted into circulation), and increased triglyceride content (Bohni *et al.* 1999; Rulifson *et al.* 2002; Ugrankar *et al.* 2015). To determine if *tsl* mutants also have such defects we performed metabolic analyses. This revealed that *tsl* null mutant (*tsl^∆^*) larvae have significantly elevated hemolymph glucose levels (Fig. 1A, *P* = 0.0002), despite consuming less food than heterozygous controls over a one hour time period (Fig. S1, *P* = 0.0040). By contrast, the concentration of circulating trehalose was unaltered in *tsl^∆^* larvae (Fig. 1B, *P* = 0.7232). In addition to the observed increase in circulating glucose, *tsl^∆^* larvae had increased triglyceride content per milligram of body weight compared to heterozygous controls (Fig. 1C, *P* = 0.0040). Taken together, the metabolic phenotype of *tsl^∆^* larvae is consistent with previous studies on *chico* and other insulin pathway mutants (Bohni *et al.* 1999; Rulifson *et al.* 2002; Ugrankar *et al.* 2015), and supports the idea that Tsl regulates the insulin signalling pathway.

**Figure 1.**
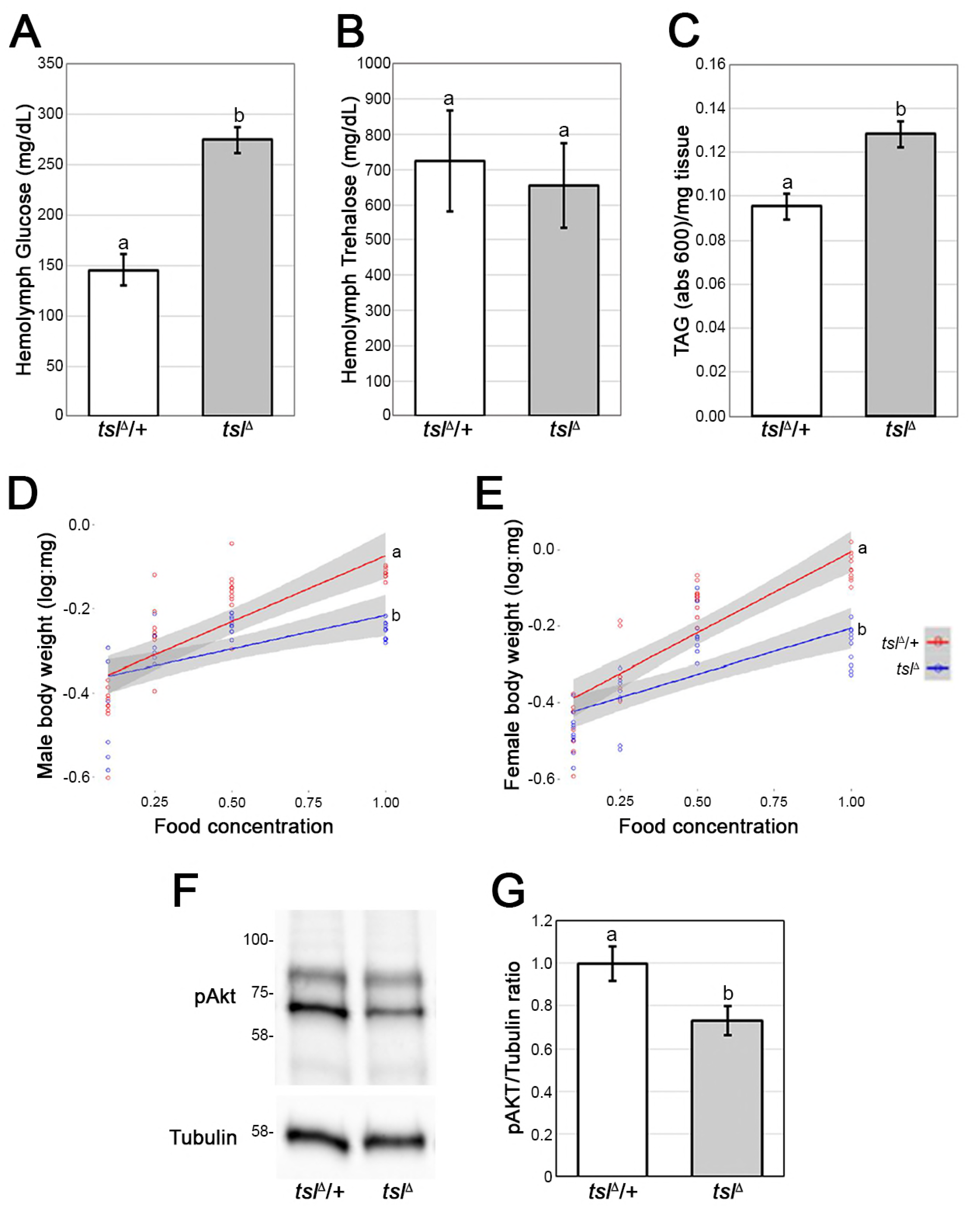
*torso-like* null mutants phenocopy mutants with reduced insulin signalling. **(A-C)** *tsl^Δ^* larvae have significantly elevated hemolymph glucose levels (A, *P* = 0.0002), unaltered hemolymph trehalose levels (B, *P* = 0.7232) and increased total triglyceride content (C, *P* = 0.0040) compared to heterozygous controls (*tsl^Δ^*/+). n = 5 groups of 10 larvae for all means. **(D, E)** The variation of both male (D) and female (E) adult body weight over different food concentrations is significantly smaller for *tsl^Δ^* animals compared to heterozygous controls (*tsl^Δ^*/+). Regression lines that differ significantly in their slopes, indicating differences in nutritional plasticity for body size between genotypes, are marked with different letters. n = 6–10 groups of at least 3 individuals for each food type. **(F)** *tsl^Δ^* larvae show reduced levels of phosphorylated Akt (pAkt). **(G)** Levels of pAkt were quantified from four biologically independent experiments, using Tubulin as a loading control. pAkt/Tubulin densities were standardised by fixing the values of *tsl^Δ^*/+ to 1. For all bar graphs, error bars represent ±1 SEM and genotypes sharing the same letter indicate that they are statistically indistinguishable from one another (*P* < 0.05, two-tailed *t* tests). The data used to generate each graph can be found in Supplementary data file 1.

Insulin signalling is also required for coupling nutrition and growth, such that body size is adjusted according to nutritional availability (Tang *et al.* 2011). For example, in wildtype flies kept under low nutritional conditions insulin signalling is downregulated, resulting in a reduced larval growth rate and decreased adult body size. Accordingly, mutants with impaired insulin signalling are unable to adjust their body size in response to nutrition to the same extent as wild type flies, as signalling is downregulated even in highly nutritious environments (Tang *et al.* 2011). To determine whether *tsl* mutants also share this feature with insulin signalling mutants we examined their adult body size when grown as larvae on foods with varying nutrient content. Diluting the larval diet to 50, 25 and 10% of control food resulted in progressively smaller adults for both male and female heterozygous controls (Fig. 1D, E). At the lowest food concentration *tsl^∆^* animals showed little difference in body size compared to heterozygous controls. However, as the food concentration increased, *tsl* mutants exhibited significantly reduced plasticity for body size (Fig. 1D, E; *P* = 0.0104 for males, *P* = 0.0038 for females; Table S1) as would be expected if insulin signalling is impaired. The reduction in plasticity for *tsl^∆^* animals was not found to be as severe as that observed in *chico* mutant animals, which appeared smaller across all food concentrations (Fig. S2, *P* < 0.0001 for both males and females; Table S2).

Next, we used immunoblotting to measure the levels of phosphorylated Akt (pAkt), a biochemical readout of insulin signalling pathway activity. Consistent with a reduction in insulin signalling, we found that *tsl^∆^* larvae had significantly lower pAkt levels compared to heterozygous controls (Fig. 1F, G, *P* = 0.0440). Taken together, these data show that many aspects of the *tsl* mutant phenotype parallel those seen in insulin signalling mutants, supporting the idea that Tsl is required for insulin signalling in response to nutrition.

To provide further evidence that Tsl acts in the insulin signaling pathway we conducted genetic interaction studies. We first asked if the delay caused by loss of *tsl* is epistatic or additive to that caused by loss of the dILPs. Consistent with previous studies (Grönke *et al.* 2010), removing dILPs 2, 3 and 5 resulted in a severe developmental delay, with a delay of ∼341 h compared to heterozygous controls (Fig. 2A, *P <* 0.0001). Loss of *tsl* alone resulted in an 18 h delay (Fig. 2A, *P <* 0.0001). In larvae mutant for *tsl* and *dilps* 2, 3 and 5 the observed developmental delay was similar to the delay seen for *dilp2-3^∆^,dilp5^3^* mutants alone (Fig. 2A, *P* = 0.2192). This result suggests that Tsl and dILPs 2, 3 and 5 act via the same signalling pathway to regulate developmental timing.

**Figure 2.**
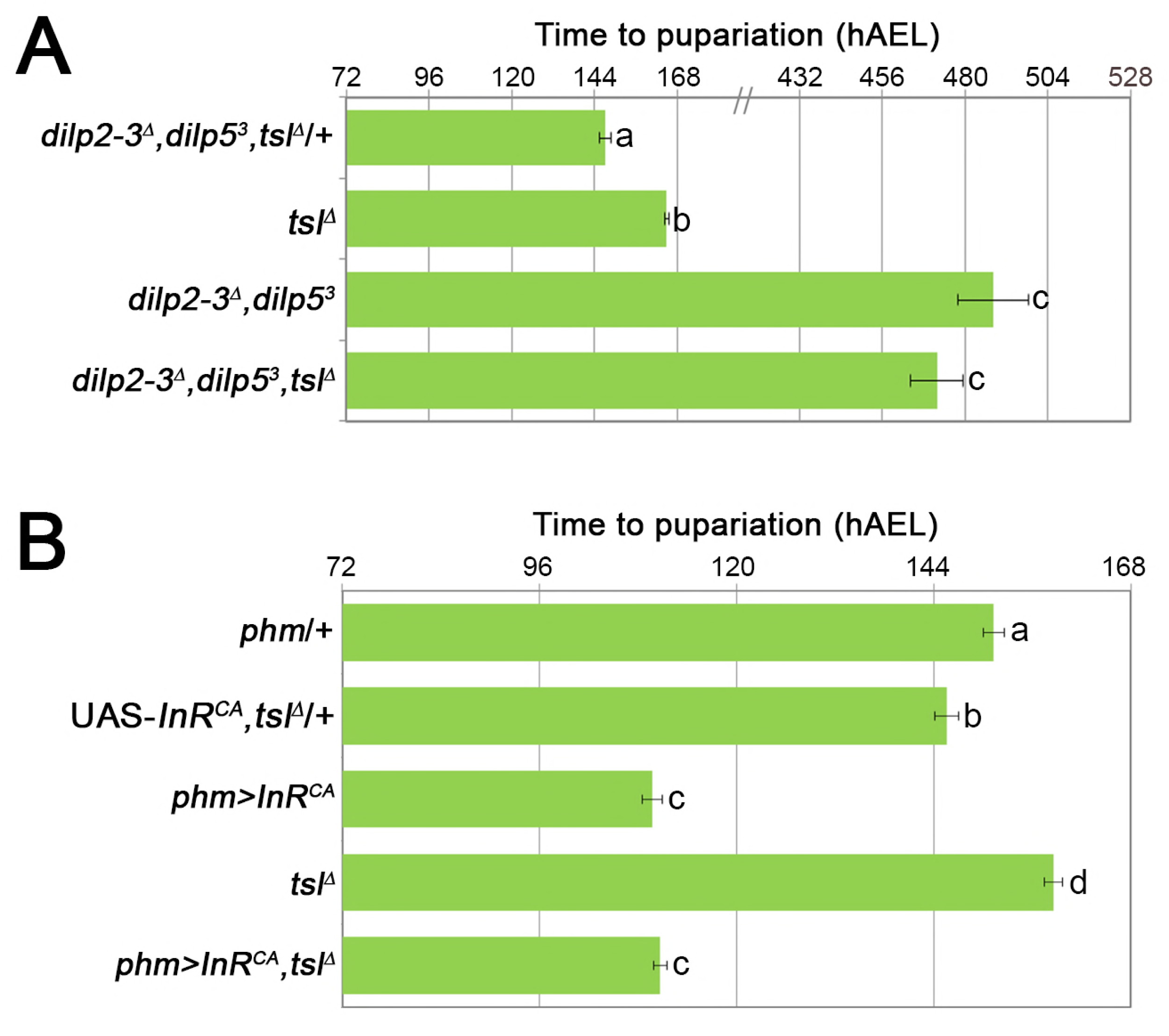
Torso-like genetically interacts with the insulin signalling pathway. **(A)** Larvae deficient for both *tsl* and *dilps 2, 3* and *5* (*dilp2-3Δ, dilp5^3^, tsl^Δ^*) show a similar delay in time to pupariation (∼325 h) to loss of *dilps 2, 3* and *5* alone (*P* = 0.2192). **(B)** The reduced time to pupariation caused by expression of *InR^CA^* in the prothoracic gland (∼42 h, *P* < 0.0001) is not suppressed by removal of *tsl* (*P* = 0.5015). h AEL = hours after egg lay. Error bars represent ±1 SEM for all graphs. Genotypes sharing the same letter indicate that they are statistically indistinguishable from one another (*P* < 0.05, ANOVA and pairwise *t* tests). n = 10 or greater for all means and no fewer than 37 individuals were tested per genotype. The data used to generate each graph can be found in Supplementary data file 1.

We next over-expressed a constitutively active and ligand-independent form of InR (*InR^CA^*) in the PG (using *phm-*Gal4) and asked whether Tsl is required for its function. We chose to manipulate InR activity specifically in the PG because it is well established as the key tissue involved in the InR-mediated regulation of ecdysone production (Mirth *et al.* 2005), and because we know that *tsl* is expressed there (Grillo *et al.* 2012). As expected (Walkiewicz & Stern 2009), over-expression of *InR^CA^* in the PG markedly reduced the time to pupariation (Fig 2B, *P* < 0.0001). This phenotype was not suppressed by loss of *tsl* (Fig. 2B, *P* = 0.5015), suggesting that Tsl activity is not required for insulin signalling downstream of InR in the PG. Taken together, the results of these two genetic interaction experiments further support the idea that Tsl acts in the insulin signalling pathway, and, if so, that it does so either upstream or at the level of InR

In the early embryo our work has led us to hypothesise that Tsl is required for the secretion of the Tor ligand, Trunk (Henstridge *et al.* 2014; Johnson *et al.* 2015). We therefore reasoned that it might regulate secretion of the dILPs, the ligands for InR. In mutants that affect dILP secretion an accumulation of dILP2 and dILP5 is observed in the insulin-producing cells (IPCs) (Geminard *et al.* 2009; Rajan & Perrimon 2012; Sano *et al.* 2015; Koyama & Mirth 2016). We therefore immunostained the IPCs in *tsl^∆^* larvae for both dILP2 and dILP5. This revealed that *tsl^∆^* larvae had a significant increase in the accumulation of both dILP2 (Fig. 3A, B; *P* = 0.0019) and dILP5 (Fig. 3C, D; *P* < 0.0001) in the IPCs compared to controls, with dILP2 accumulating to a lesser extent than dILP5.

**Figure 3.**
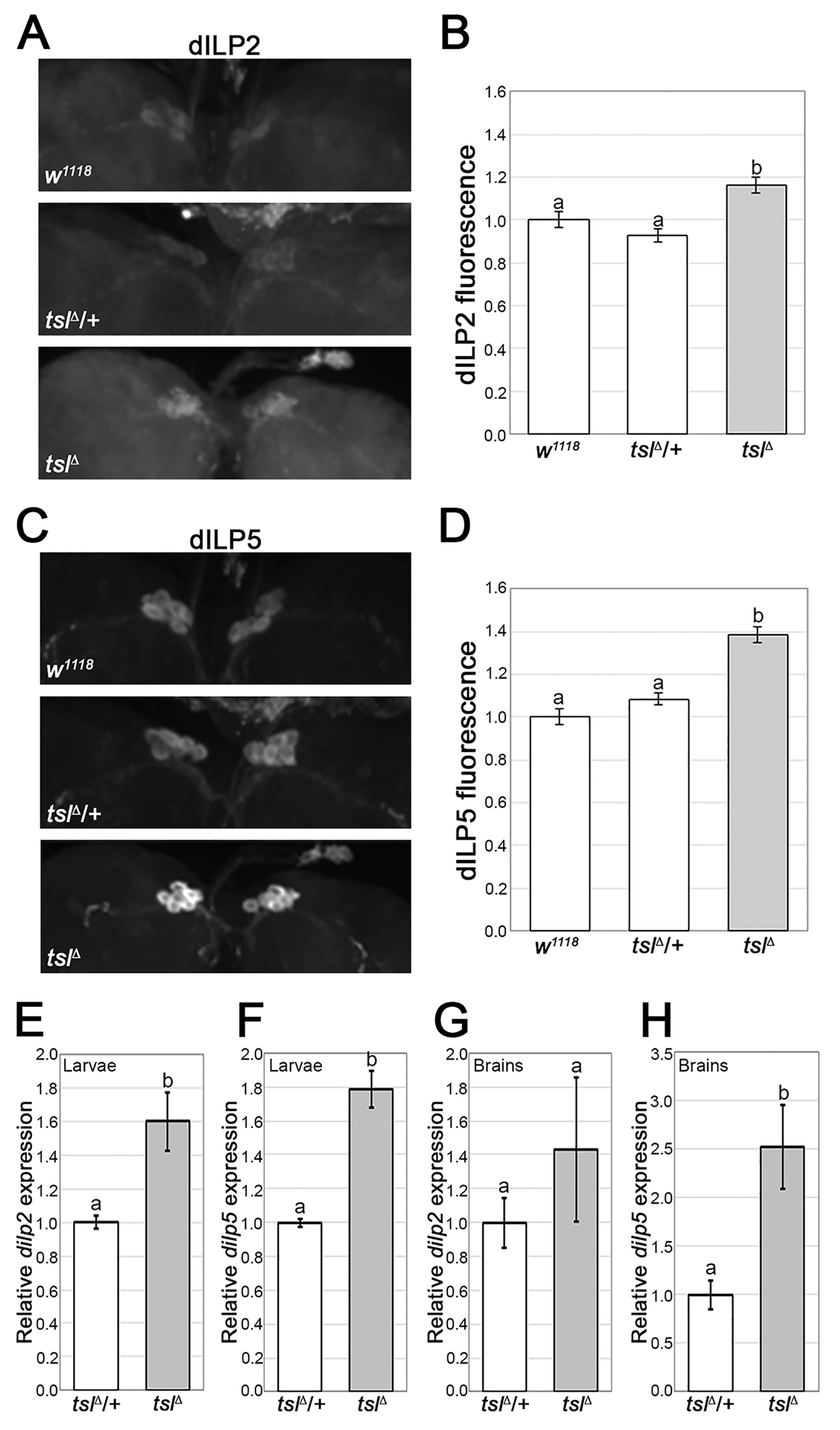
*torso-like* mutants show increased insulin-like peptide expression and accumulation in the insulin-producing cells. **(A-D)** Removal of *tsl* increases the accumulation of dILP2 (A, B) and dILP5 (C, D) in the insulin-producing cells (IPCs). dILP protein levels were standardised by fixing the values of *w^1118^* to 1. n = 35 or greater for all means. **(E, F)** Expression of *dilp2* (E, *P* = 0.0240) and *dilp5* (F, *P* = 0.0016) is significantly increased in *tsl^Δ^* larvae compared to controls (*tsl^Δ^*/+). **(G, H)** *dilp2* expression in the larval brain is not significantly altered in *tsl* mutants (G, *P* = 0.4243), however expression of *dilp5* is significantly elevated (H, *P* = 0.0009). Expression levels were normalised using an internal control, Rp49, and then standardised by fixing the values of *tsl^Δ^*/+ larvae to 1. n = 3–5 for all means and no fewer than 75 individuals were tested per genotype. For all graphs, error bars represent ±1 SEM and genotypes sharing the same letter indicate that they are statistically indistinguishable from one another (*P* < 0.05, ANOVA and pairwise *t* tests). The data used to generate each graph can be found in Supplementary data file 1.

While the observed accumulation of dILP2 and dILP5 could reflect a defect in their secretion, it was also possible that the observed accumulation reflects elevated expression of these peptides. Increased *dilp2/5* expression is commonly observed when there is a systemic reduction in insulin signalling caused by insulin resistance in peripheral tissues (Musselman *et al.* 2011; Pasco & Leopold, 2012). To determine whether the observed accumulation of dILP2 and dILP5 results from their elevated expression we quantified *dilp2* and *dilp5* mRNA levels. This showed that the expression of both *dilp2* (Fig. 3E, *P* = 0.0240) and *dilp5* (Fig. 3F, *P* = 0.0016) was elevated in *tsl^∆^* larvae compared to heterozygous controls. As low levels of *dilp* expression have previously been detected in larval tissues other than the IPCs (Brogiolo *et al.* 2001), we also quantified *dilp2* and *dilp5* mRNA levels specifically in the larval brain. We found no significant difference in the expression of *dilp2* (Fig. 3G, *P* = 0.4243) in *tsl^∆^* brains compared to heterozygous controls, however expression of *dilp5* was significantly elevated (Fig. 3H, *P* = 0.0009). Taken together, these findings suggest that Tsl influences *dilp* expression during larval development.

We next asked if the role of Tsl in insulin signalling is due to a function in the PG. Previously, Grillo *et al.* (2012) showed that *tsl* is expressed in the PG and that RNAi knockdown of *tsl* specifically in this tissue results in a significant developmental delay. While we were unable to reproduce this result using the publically available RNAi lines (Johnson *et al.* 2013), when we used the same RNAi line used in the Grillo *et al.* study (originally generated by Furriols *et al.* 2007) we did observe a developmental delay phenotype (Fig. 4A, *P* = 0.0018). By contrast, no phenotype was observed when we knocked down *tsl* expression specifically in the fat body using *c7*-Gal4 (Fig. 4B, *P* = 0.9201).

**Figure 4.**
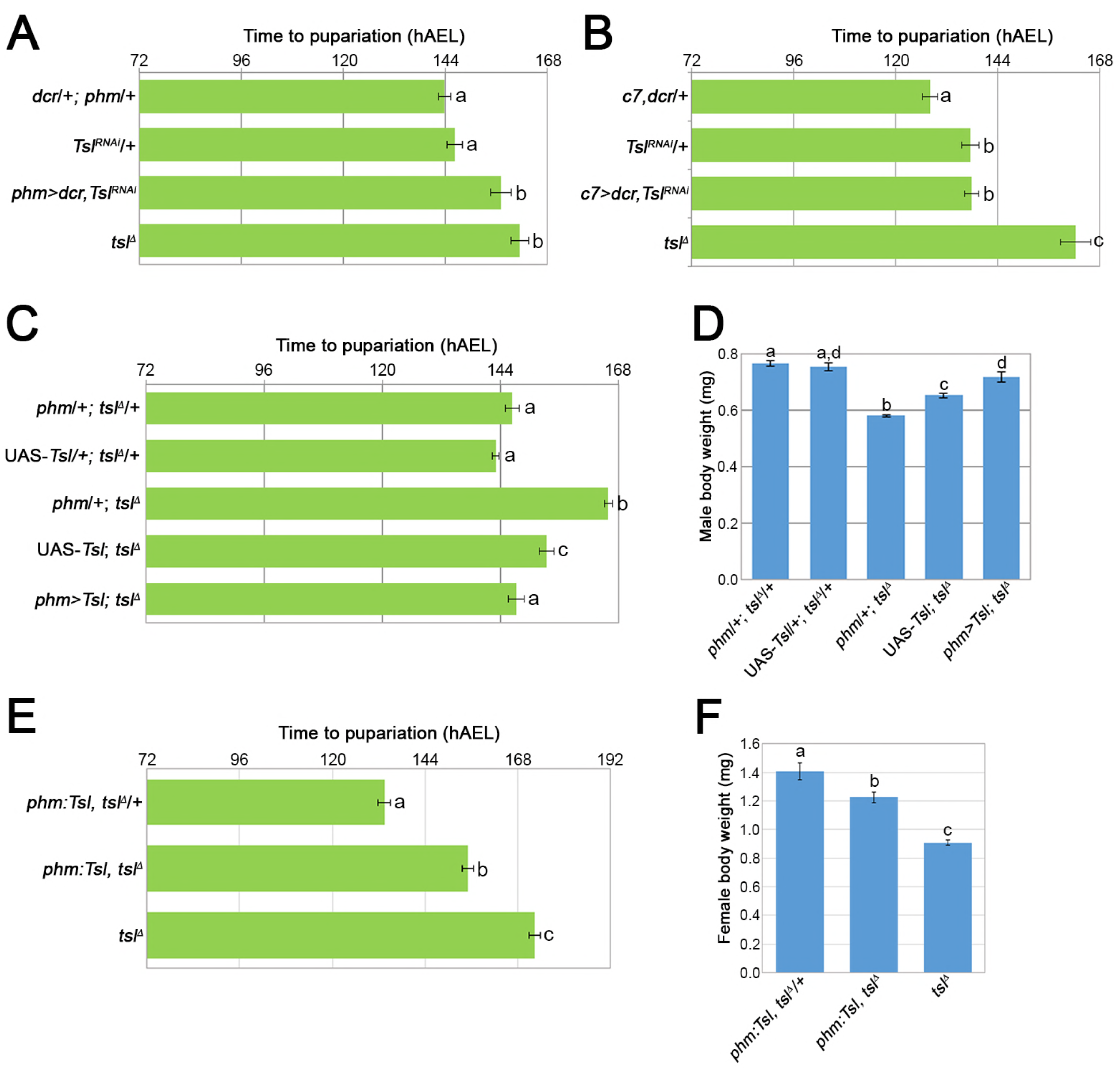
Torso-like is required in the prothoracic gland to regulate both developmental timing and body size. **(A)** Knockdown of *tsl* specifically in the prothoracic gland (using *phm-*Gal4) results in a significant developmental delay (∼11 h, *P* = 0.0018) that is similar to the delay observed for *tsl^Δ^*. **(B)** No developmental delay is observed when *tsl* is knocked down specifically in the fat body (using *c7-*Gal4). **(C, D)** Expression of a UAS-*tsl:RFP* (UAS*-tsl*) transgene specifically in the prothoracic gland (using *phm-*Gal4) rescues both the developmental delay (C) and reduced adult body size (D) of *tsl^∆^* homozygotes (*P* < 0.0001 for both delay and body size compared to *tsl^∆^*). **(E, F)** The developmental delay (C) and reduced adult body size (D) or *tsl* mutants is partially rescued by the *phm:Tsl:3Myc:eGFP* (*phm:Tsl*) construct (*P* < 0.0001 for both delay and body size compared to *tsl^∆^*). h AEL = hours after egg lay. Error bars represent ±1 SEM for all graphs. Genotypes sharing the same letter indicate that they are statistically indistinguishable from one another (*P* < 0.05, ANOVA and pairwise *t* tests). n = 10 or greater for all means and no fewer than 37 individuals were tested per genotype. The data used to generate each graph can be found in Supplementary data file 1.

We also asked if PG-specific expression of *tsl* could rescue the growth defects of *tsl* mutants. When we expressed a UAS-*Tsl:RFP* transgene (UAS-*Tsl*) in the PG using *phm-*Gal4 we found this completely rescued both the developmental delay (Fig. 4C, *P* < 0.0001) and small body size (Fig. 4D, *P* < 0.0001) of *tsl* mutants. However, this transgene also partially rescued the *tsl^∆^* phenotypes in the absence of the Gal4 driver, most likely due to leaky transgene expression. To overcome this problem, we generated a genomic rescue construct in which the *phm* promoter sequence (from *phm-*Gal4) was fused to the *tsl* coding sequence C-terminally tagged with three tandem Myc epitopes and the eGFP coding sequence (*phm:Tsl*). We found that this transgene partially rescued both the developmental delay (Fig. 4E, *P* < 0.0001) and small body size (Fig. 4F, *P* < 0.0001) of *tsl* null mutants. Together with the RNAi experiments, these data strongly suggest that *tsl* expression is required in the PG for regulating developmental timing and body size. However, we are unable to rule out the possibility that *tsl* is also required in other tissues for these roles.

How might Tsl function in the PG to regulate systemic insulin signalling? Because Tsl is a secreted protein this could be explained if Tsl is secreted from the PG into the hemolymph. We therefore asked if Tsl is found in the larval hemolymph. To do this we used a genomic rescue construct that carries ∼3kb of promoter and the *tsl* coding sequence N-terminally tagged with three tandem hemagglutinin (HA) epitopes (*gHA:Tsl*; Jimenez *et al.* 2002). This construct has previously been shown to completely rescue the developmental delay and reduced body size of *tsl*^∆^ animals (Johnson *et al.* 2013). Using immunoblotting, we were able to clearly detect *gHA:Tsl* in the larval hemolymph (Fig. 5A). We further asked if PG-produced Tsl enters the hemolymph by expressing a functional C-terminally tagged Tsl transgene (UAS-*Tsl:HA*) in the PG (*phm-*Gal4) and performing western blots on protein extracted from larval hemolymph. This revealed that *Tsl:HA* protein was present in the hemolymph (Fig. 5B). While this is an over-expression situation, we reason that because Tsl is endogenously expressed in the PG it is likely that at least a proportion of the total Tsl in circulation originates from the PG. However, it remains possible that the Tsl we detect in the hemolymph with the genomic construct is produced and secreted from another tissue.

**Figure 5.**
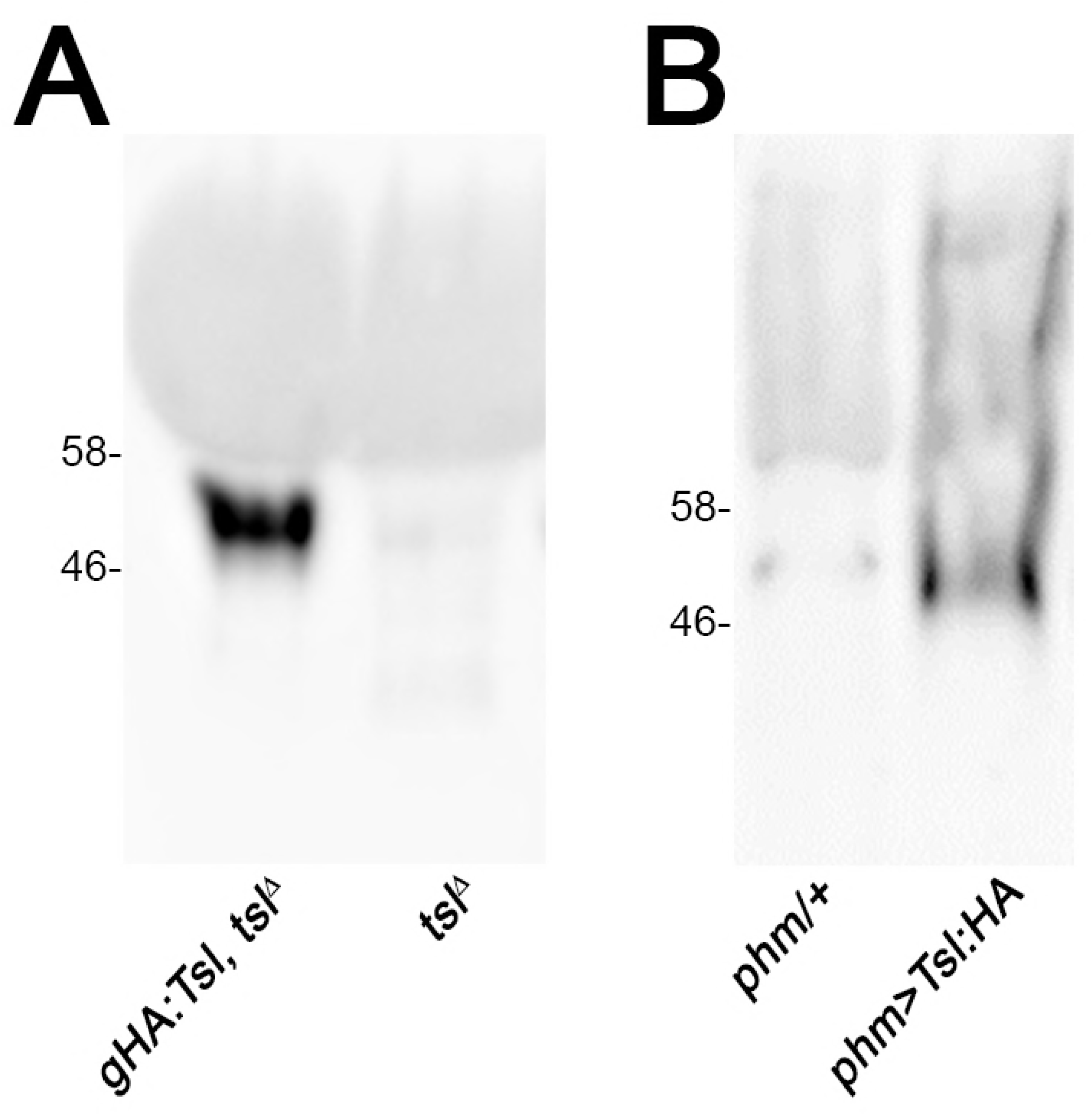
Torso-like is secreted from the prothoracic gland into the larval hemolymph. **(A**) Immunoblot (anti-HA) of Tsl expression in the larval hemolymph. **(B)** Tsl is also detected in the hemolymph via immunoblotting (anti-HA) when expressed in the prothoracic gland using *phm-*Gal4.

## DISCUSSION

Our data presented here provide compelling evidence that Tsl is secreted into the hemolymph and regulates growth and developmental timing via the insulin signalling pathway. While the *tsl* mutant phenotypes described here closely resemble those observed when insulin signalling is reduced in the entire organism, it should be noted that loss of *tsl* has a less severe effect on the pathway compared to mutations in other genes. For example, loss of function mutations in *InR* are homozygous lethal and only a few heteroallelic combinations produce viable adults in which growth defects can be observed (Chen *et al.* 1996). By comparison, mutations in the adaptor protein Chico do not result in lethality but rather cause severe growth and metabolic defects (Bohni *et al.* 1999). Here we find that the defect in nutritional plasticity for body size is not as severe in *tsl* mutants as it is in *chico* mutants. Our findings therefore suggest that Tsl regulates, but is not essential for insulin signalling.

How might Tsl regulate insulin signalling throughout the body? One possibility is that Tsl regulates the insulin response in all tissues by acting in conjunction with InR (Fig. 6A). Loss of insulin response in *tsl* mutants could then result in the increased *dilp* expression that we observe. Alternatively, Tsl may act to influence the activity of the dILPs, which in turn regulate systemic insulin signalling. There are two main tissues that are known to regulate dILP activity (for a complete review of the regulation of dILP production and secretion see Nassel & Vanden Broeck 2016). One is the fat body, which produces and releases important regulators of dILP secretion in response to intracellular nutrient levels (Fig. 6B; Geminard *et al.* 2009; Rajan & Perrimon, 2012; Sano *et al.* 2015; Delanoue *et al.* 2016; Koyama & Mirth 2016). Interestingly, a recent study found that knocking down *tor* specifically in the fat body led to a decreased body size, leading the authors to suggest that Tor acts in the fat body to influence insulin signalling via an unknown mechanism (Jun *et al.* 2016). Given the key role of Tsl in the regulation of Tor activity during early embryogenesis, it is therefore possible that Tsl acts in the Tor pathway in the fat body. However, fat body-specific knockdown of *tor* does not result in a developmental delay (Jun *et al.* 2016), thus this would seem unlikely to be the only role for Tsl in regulating growth and developmental transitions. In addition, experiments that we have performed to detect Tsl or knockdown its expression in the fat body have not provided any evidence of Tsl expression or function in this tissue. Determining if Tsl acts in the fat body to regulate dILP activity, either with Tor or with another pathway, thus requires further investigation.

**Figure 6.**
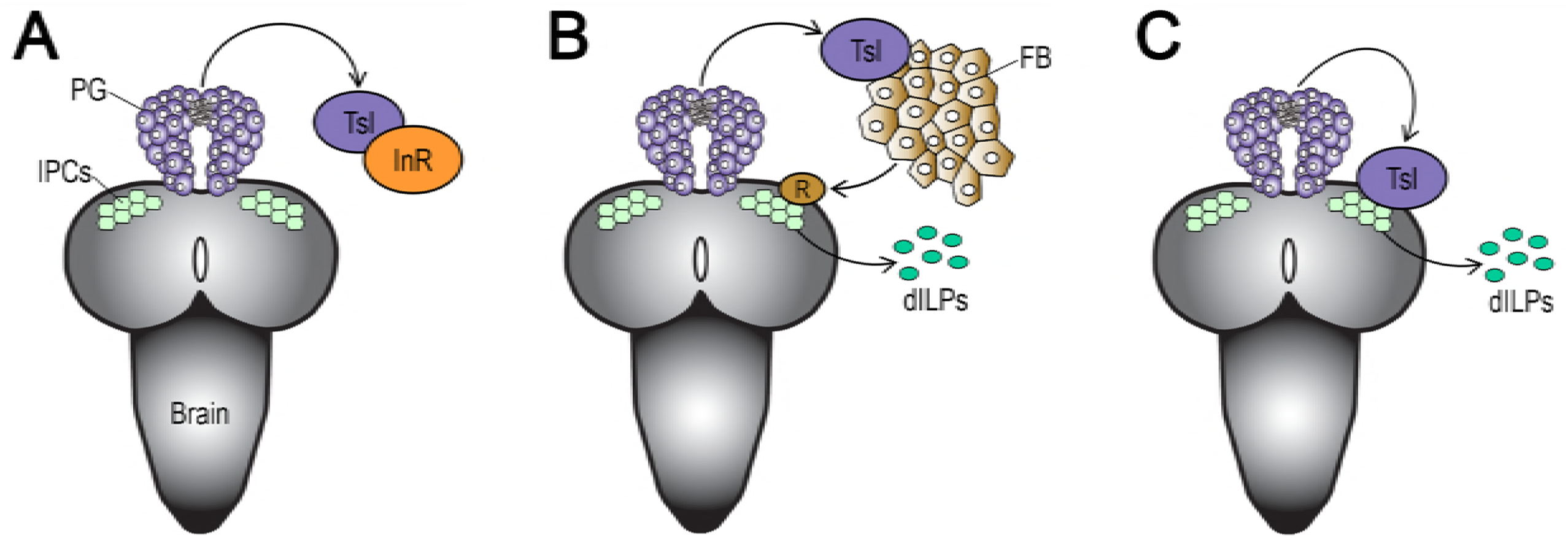
Possible mechanisms of Torso-like function from the prothoracic gland. **(A)** Tsl is secreted from the prothoracic gland (PG) into circulation where it acts in conjunction with InR in all tissues to modulate insulin signalling. **(B, C)** Tsl influences the activity of the dILPs in the insulin-producing cells (IPCs), which in turn regulate systemic insulin signalling. In this model, Tsl could act on either (B) the fat body (FB), which is responsible for producing and releasing important regulators of dILP secretion (R, brown oval) such as CCHamide-2, Unpaired-2, Growth-blocking peptides 1 and 2 and Stunted, or (C) the IPCs to directly control dILP activity.

An alternative possibility is that Tsl acts directly on the IPCs to control the activity of the dILPs (Fig. 6C). Given the close proximity of the IPCs to the PG, this perhaps fits better with the known role of Tsl in the early embryo, where it is secreted from the follicle cells and acts locally on Tor signalling (Jimenez *et al.* 2002; Stevens *et al.* 2003). It is therefore possible that the systemic effects of Tsl are due to a role in regulating dILP expression and/or secretion in the IPCs. These ideas could be tested in future by experiments such as examining the kinetics of dILP secretion in *tsl* mutants following starvation, or testing if artificially stimulating dILP release can rescue the *tsl* mutant phenotype. Understanding the exact role of Tsl in this system will provide fundamental insights into the mechanisms that regulate the evolutionarily conserved insulin signalling pathway, as well as the role of MACPF proteins in developmental signalling events.

## ACKNOWLEDGEMENTS

We thank Karyn Moore, Lauren Forbes Beadle, Katherine Shaw and the Australian *Drosophila* Biomedical Research Facility (OzDros) for technical support; Jordi Casanova and Michael O’Connor for providing fly stocks; and Pierre Léopold for the dILP2 and dILP5 antibodies. M.A.H is a National Health and Medical Research Council (NHMRC) Early Career Fellow. This work was supported by an Australian Research Council (ARC) grant to C.G.W, C.K.M and T.T.

## AUTHOR CONTRIBUTIONS

C.G.W conceived the experiments, interpreted the data and led the work. M.A.H conceived the experiments, performed the experiments and interpreted the data. L.A, T.K and T.K.J performed experiments. J.C.W, T.T and C.K.M interpreted the data. M.A.H and C.G.W wrote the paper with assistance from all authors.

